# EMG-to-torque models for exoskeleton assistance: a framework for the evaluation of *in situ* calibration

**DOI:** 10.1101/2024.01.11.575155

**Authors:** Lucas Quesada, Dorian Verdel, Olivier Bruneau, Bastien Berret, Michel-Ange Amorim, Nicolas Vignais

## Abstract

In the field of robotic exoskeleton control, it is critical to accurately predict the intention of the user. While surface electromyography (EMG) holds the potential for such precision, current limitations arise from the absence of robust EMG-to-torque model calibration procedures and a universally accepted model. This paper introduces a practical framework for calibrating and evaluating EMG-to-torque models, accompanied by a novel nonlinear model. The framework includes an *in situ* procedure that involves generating calibration trajectories and subsequently evaluating them using standardized criteria. A comprehensive assessment on a dataset with 17 participants, encompassing single-joint and multi-joint conditions, suggests that the novel model outperforms the others in terms of accuracy while conserving computational efficiency. This contribution introduces an efficient model and establishes a versatile framework for EMG-to-torque model calibration and evaluation, complemented by a dataset made available. This further lays the groundwork for future advancements in EMG-based exoskeleton control and human intent detection. This work has been submitted to the IEEE for possible publication. Copyright may be transferred without notice, after which this version may no longer be accessible.

## I Introduction

**A**CTIVE exoskeletons are a major potential tool to support neurorehabilitation protocols and prevent the appearance of musculoskeletal disorders (MSDs) at work. Concerning rehabilitation, they can be used to improve gait and upper limb control during daily activities for those who have suffered a stroke or a spinal cord injury [1]–[4]. Exoskeletons can also be used to decrease the biomechanical load on the body [5], thus contributing to preventing the onset of MSDs when the workspace cannot be adapted to improve ergonomics [6], although more evidence is needed [7], [8]. In such situations, integrating the human intention into the exoskeleton’s control loop is necessary to ensure safe, appropriate, and intuitive assistance [9].

Human intention detection (HID) involves continuously estimating the motion of one or more human joints. There are several approaches to this aim. Kinematics can be used to anticipate the trajectory of a movement based on prior knowledge of typical trajectories. These techniques, often based on probabilistic models [10], [11], have already been used to assist with an active exoskeleton [12]. Another method is to employ a bio-electric signal and a model to estimate the movement. Data sources such as mechanomyography (MMG), force myography (FMG) [13], electromyography (EMG), or a fusion of signals [14], [15] may be processed to control an exoskeleton. Thanks to the electromechanical delay, *i.e*., the latency between EMG signal and muscle force production [16], human intention detection can be predictive: the movement estimation can occur before the onset of the movement itself. This potential predictive capability can be advantageously used in real-time assistive systems, making EMG signals an appealing choice for human/machine interactions [17]–[19].

Several approaches can be employed to predict intent based on EMG signals, but two general categories can be defined: i) categorical classification and ii) continuous regression. Movement classification tends to use convolutional neural networks [20], [21], support vector machine [22], [23], gaussian mixture models [24] and other various techniques [25] linked to pattern recognition. This type of HID allows to predict which type of movement is performed (*e.g*., elbow flexion/extension) but has to be trained on a predetermined set, preventing its generalization. On the other hand, continuous movement regression aims to predict a continuous feature of the movement, such as joint angle or torque. This is generally achieved through a mathematical model linking EMG signals to the predicted feature. For example, neuromusculoskeletal models [26]–[simple linear regressions [17], [30]–[various synergy-based [31], [33]– and other custom models [37]– currently coexist. However, comparing models in the literature is difficult due to methodological differences in evaluation. To provide accurate predictions, continuous regression typically requires first calibrating the model to fit its internal parameters to an individual [29]. Some studies propose to estimate joint torques from EMG signals during isometric tasks [29], [31], [32], [40], [42], and specific or periodic dynamic tasks [28], [30], [33], [39], [43]. However, similarly to categorical classification, a realistic general assistive system cannot be restricted to isometric or predetermined tasks. Individualizing a calibration procedure might also involve a long computation time or specialized equipment, reducing its practicality for future commercial or clinical applications. Therefore, developing a model that allows short *in situ* calibration procedures capable of performing HID for arbitrary movements is crucial. To validate and compare models, the data on which the model is evaluated must be distinct from the data used for individualized calibration (*e.g*., different movements, efforts, etc.). Apart from very few studies using this type of cross-validation [32], most do not exploit this aspect, thus increasing the risk of overfitting models and misinterpreting their performance.

In the present paper, we address these concerns by: i) introducing an *in situ* calibration framework enabling a generic calibration of upper limb EMG-to-torque models, ii) applying this framework to elbow and shoulder flexion/extension concurrently and separately, iii) evaluating and comparing the performances obtained by four models, one of which is introduced. This paper is structured as follows: first, we provide an overview of the experimental protocol that produced our dataset, including the exoskeleton control law, arbitrary task generation, and EMG placement; next, we explain the data processing and model evaluation procedures; then, we describe the four EMG-to-torque models we evaluated; and finally, we present our results, along with their limitations and potential future directions.

## II. Methods

### A. Participants

17 healthy subjects (11 males, age 28.2 ± 7 years, height 175.4 ± 7 cm, weight 70 ± 11 kg) took part in the experiment. The experimental protocol was approved by the ethics committee of Université Paris-Saclay (CER-PS-2021-048/A1). Participants signed a written informed consent form before starting the experiment.

### B. Material

#### 1) Exoskeleton

The ABLE upper limb active exoskeleton [45] was used to impose a viscous resistive force on the participant’s wrist to induce torques in its elbow and shoulder.

Its movement was limited to a parasagittal plane, only allowing flexion and extension of the elbow and the shoulder. Participants were attached to the exoskeleton through a previously developed mechanical interface [44] composed of an orthosis fixed on a force/torque (FT) sensor (1010 Digital FT, ATI, Apex, USA) itself attached to a slider on the exoskeleton’s forearm. This mechanical interface includes a passive rotation between the exoskeleton and the human limb, which reduces unwanted hyperstatic constraints and increases comfort while permitting the measurement of interaction forces [46].

#### 2) Electromyography

Electromyographic signals were recorded using wireless MiniWave sensors with a WavePlus receiver (Cometa, Bareggio, Italy). The receiver was connected via USB to a computer, and the acquisition was performed through the Qualisys Track Manager (QTM) software (Qualisys, Göteborg, Sweden). Muscle head recording locations were identified following SENIAM recommendations [47]. The skin was locally shaved, cleaned with alcohol pads, and dried before placing the Ag/AgCl EMG electrodes (F3010, Fiab, Firenze, Italy). The muscle set included the long and short head of the biceps (BICShort, BICLong), long, lateral, and median heads of the triceps (TRILong, TRILat, TRIMed), brachioradialis (BRD), brachialis (BRA), posterior, medial, and anterior heads of the deltoids (DELTPost, DELTMed, DELTAnt), clavicular head of pectoralis major (PECT), and latissimus dorsi (LATI). The detailed placement of EMG sensors on the muscles is provided in supplementary material S2.

#### 3) Motion capture

Kinematic data of the participant and the exoskeleton were captured using a total of 10 Oqus 500+ motion capture cameras (Qualisys, Göteborg, Sweden). Eight 10 mm markers were placed on anatomical landmarks, including the acromion, seventh cervical vertebra, sternal end of the clavicle, styloid processes of the ulna and radius, lateral and medial epicondyles and distal end of the second metacarpal bone (see supplementary S1.a). Additionally, two markers were placed on the anterior and posterior sides of the arm, specifically at the midpoint of the upper arm. Additional markers were placed on the exoskeleton and orthosis (see Fig. 1 and supplementary S1.b) to track their movement for later data synchronization and frame transformation purposes. Motion capture data was recorded in synchronization with EMG using the QTM software.

**Fig. 1:**
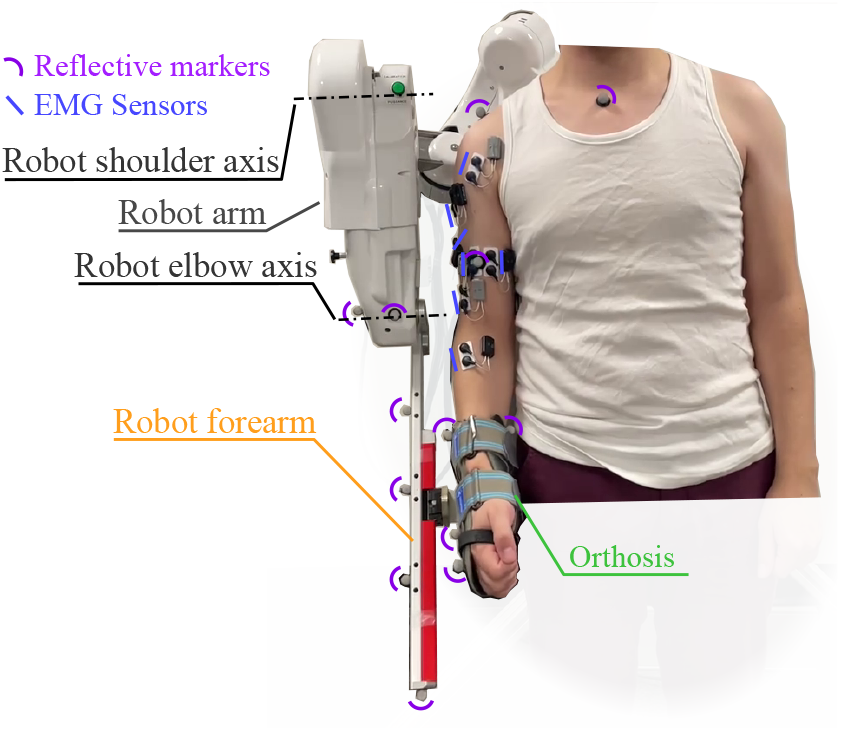
Participant with EMG sensors and reflective markers in the exoskeleton. Visible reflective markers and EMG sensors are highlighted with pink and blue lines, respectively. The setup corresponds to the multi-joint condition.

### C. Procedure

#### 1) MVC and anthropometric measurements

Participants first completed maximum voluntary contraction (MVC) tasks for flexion and extension of the elbow and shoulder. For the elbow, they placed their arm vertically against their body, resting their wrist on a horizontal bar set at the same height as their elbow joint. To perform flexion and extension MVCs, they adjusted the position of their wrist either above or below the bar and pushed against it. The same process was applied to the shoulder joint by moving the bar to the shoulder level and asking the participant to extend their elbow fully. Each task was repeated twice and lasted for 3 seconds. Following the MVC trials, after placing markers on specific anatomical landmarks as described in Section II-B3, participants were instructed to stand still within the motion capture area to capture a frame of a static pose.

#### 2) Motor tasks

The experiments were based on a trajectory tracking task and divided into two sessions. During the first session, participants tracked a trajectory using only elbow flexion/extension (*i.e*., single-joint). During the second session, participants tracked a trajectory using elbow and shoulder flexion/extension (*i.e*., multi-joint). Two different kinematic arrangements of the exoskeleton were employed, depending on the condition (see Fig. 2.a). The first setup enabled only the elbow to rotate while the wrist was secured to a slider. The second configuration allowed both the elbow and the shoulder to rotate. As previously stated in the material section, these two configurations avoided hyperstaticity between the subject and the exoskeleton, reducing unwanted interaction efforts. The trajectory was projected on a screen positioned frontally for the single-joint task and sagittally for the multijoint task (see Fig. 2.b). Visual feedback included a yellow dot representing the participant’s current position, synchronized with white dots depicting the desired trajectory scrolling across the screen. For multi-joint tasks, red dots indicated specific positions participants were expected to reach at a given time. Participants earned a point each time the yellow dot coincided with a target dot, displaying an interactive score to enhance engagement. A video of the experiments in single-joint and multi-joint conditions can be viewed in supplementary material S3 and S4.

**Fig. 2:**
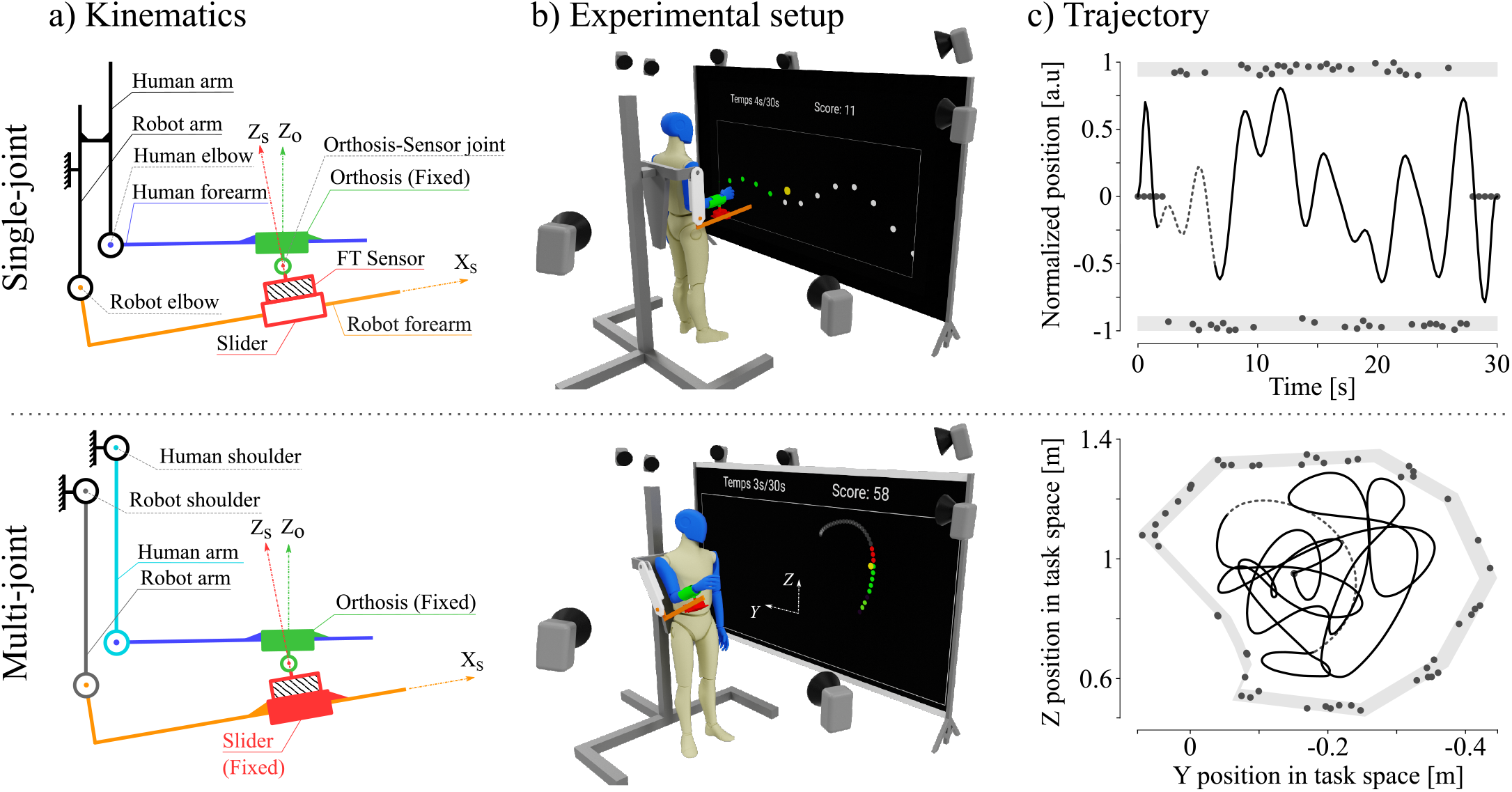
Illustration of the different experimental conditions. **a)** During the single-joint task, the movement was limited to elbow flexion/extension. To limit hyperstatic constraints in the human-exoskeleton kinematic chain and increase comfort, a passive translation along *X*_*s*_ (slider) and a passive rotation around *Y*_*s*_ were allowed by the physical interface [44]. Both shoulder and elbow flexion/extension were involved during the multi-joint task. Participants were only attached to the exoskeleton at the wrist level, which implied that the slider needed to be fixed to transmit efforts from the human to the exoskeleton. The *Z*_*s*_ axis represents the sensor frame, while the *Z*_*o*_ axis represents the orthosis frame. It is necessary to consider this angular misalignment to compute accurate inverse dynamics. **b)** Participants were positioned in front of a projected screen during the tasks. The trajectory that needed to be tracked was visualized as white dots, while a yellow dot represented the participant’s current position. As participants successfully touched the white dots, they would turn green, indicating a successful catch. The displayed score corresponded to the number of dots participants successfully caught. In the single-joint task, the height of the yellow dot representing the participant’s position was determined by the angular position of their elbow. During the multi-joint task, participants were turned 90 degrees, and the Cartesian coordinates of their wrist determined the position of the yellow dot. Two screen dimensions were effectively utilized for spatial representation, where white dots represented the future trajectory, and red dots represented the set of accessible positions during the current period (500 *ms*). **c)** In both cases, B-splines are being used to produce the trajectories with control points (gray dots) generated into the light-gray area that contains the reachable workspace. Trajectory examples are shown as black lines. The section of the trajectory currently displayed on the screen of the *b* panel is dashed.

### a) Exoskeleton Control

The exoskeleton was controlled to impose a viscous resistance at the elbow and shoulder joints, increasing the muscle effort induced by the task. This allowed to extend the range of measured muscle activities and, therefore, the range of validity of the resulting calibration. Furthermore, to reduce the effort asymmetry generated by the exoskeleton’s weight between upward and downward movements, it compensated for its weight based on a former identification [48]. For the single-joint condition, the resulting exoskeleton control torque *τ*_*e*_ was computed as follows:

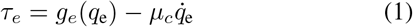

with ***µ***_*c*_ representing the viscous coefficient, and 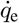 denoting the angular velocity of the exoskeleton’s elbow joint. For the multi-joint condition, the same gravity compensation and viscous resistance scheme was applied, yielding control torques computed from task space coordinates using the following equation:

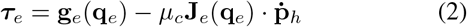

where **J**_*e*_(**q**_*e*_) and **q**_*e*_ respectively represent the Jacobian matrix and the angular positions of the exoskeleton and 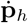 the velocity vector of the hand expressed in the task space. In both equations, *g*_*e*_ or **g**_*e*_ denotes the gravity model of the exoskeleton.

### b) Trajectory generation

The target trajectories were generated using random b-splines (see Fig. 2.c). A set of random control points was generated in time and space to create a candidate trajectory. Among the generated trajectories, the one maximizing the mechanical work while covering the whole reachable workspace was chosen. This approach to random trajectory generation can produce a broad spectrum of trajectories, ranging from flat to very curved. To effectively scan the workspace, two criteria were established to select trajectories that displayed a wide range of movements. These criteria were the amount of mechanical work and the evenness of its distribution in the workspace. For each component *I* of the trajectory (*e.g*., *i* ∈ [*Y, Z*] for multi-joint condition), according to equations 1 and 2, a substitute to the mechanical work created by the viscous field can be computed as

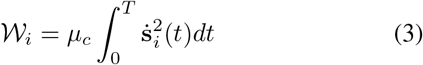

where *T* was the duration of the trajectory, and **s** corresponds to *q*_*e*_ and **p**_*h*_ for single and multi-joint conditions, respectively. The distribution of mechanical work within the movement can be assessed by analyzing partial work, which quantifies the amount of work dedicated to movement in either the positive or negative direction:

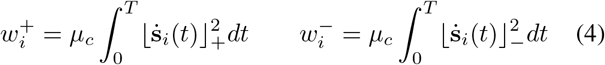

where 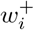 and 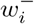 respectively denote the work of the movement in the upward (*i.e*., positive and denoted 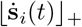) and downward (*i.e*., negative and denoted 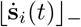) directions. The normalized work balance criterion is then expressed as the mean of relative partial work contributions. With the total work as the sum of individual components work 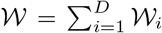, the criterion is thus expressed as:

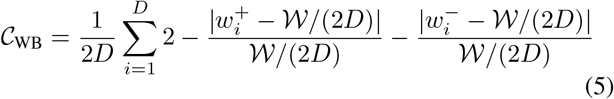

here, *D* represents the dimension of the task space (*e.g*., *D* = 1, 2 for single and multi-joint conditions, respectively). For example, in multi-joint condition, forward, backward, upward, and downward movements should all account for a quarter of the total work. With equation 5, trajectories with an evenly distributed work throughout space maximize 𝒞_WB_. Before executing the task, a hundred trajectories were generated, and their respective scores were computed as

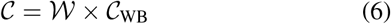

Finally, the trajectory with the highest score is selected and used for the task.

#### 3) Trials

Before initiating the main trials, preliminary test tasks were conducted to iteratively set the viscous coefficient for the robot controller. In these initial trials, a baseline viscous coefficient was employed, and participants were requested to rate their perceived effort on a subjective Borg scale. Based on the participants’ ratings, the viscous coefficient was then manually adjusted to achieve a moderate effort. Then, twenty trials were performed, each lasting thirty seconds and featuring a different random trajectory. Ten trials were conducted for each condition: single-joint and multi-joint. The trial procedure was as follows: a random trajectory was generated, and participants were instructed to stand ready. A countdown with an accompanying sound cue was initiated when the trial began. A score, displayed on the top right of the screen, kept track of the number of successfully caught dots, encouraging participants to track the trajectory accurately. A one-minute rest period was observed between each trial to prevent muscle fatigue.

### D. Data processing

#### 1) Data synchronization

EMG and motion capture data were synchronized using the QTM software. However, the exoskeleton kinematic data obtained from internal encoders and force/torque measurements were recorded separately, necessitating temporal synchronization with the EMG and motion capture data. To address this, the exoskeleton’s positions were measured using its internal encoders and the motion capture system. The alignment process consisted of determining the optimal temporal shift that minimized the root mean square error between the encoder-informed and motion-capture-informed positions of the exoskeleton arm. Once this optimal shift was determined, it was applied to the force/torque data to ensure proper alignment across all data sources.

#### 2) EMG feature extraction

EMG signals underwent a series of filtering stages. Initially, to eliminate noise from movement artifacts and external electromagnetic disturbances, a fourth-order Butterworth bandpass filter with a frequency range of 20-450Hz was utilized. Following this, the signal was rectified. Subsequently, to obtain useful information from the signal, the EMG envelope was extracted using a 3Hz fourth-order Butterworth low-pass filter [49]. These filtering operations were conducted using a forward-backward method to prevent any phase shift. The same filtering technique was also applied to the EMG data from MVC tests, during which the peak values were computed for each participant and muscle involved. Lastly, the EMG signals were normalized based on these peak values.

#### 3) Anthropometrics

Data collected during the static motion capture phase was used to scale an upper limb model [50] using OpenSim’s scaling tool [51]. This process involved adjusting the dimensions of the model’s bones to match the marker spacing on anatomical landmarks. By adapting the bone dimensions, the model accurately represents the proportions of the individual’s limb. Furthermore, the limb masses were computed by considering the ratio of the new bone dimensions to the original dimensions of Holzbaur’s model. This ensured that the model incorporates the appropriate mass distribution (mass matrix) to simulate the upper limb dynamics accurately.

#### 4) Kinematics

Although human kinematics can be deduced from the exoskeleton’s internal encoders for practical scenarios, it was instead computed from motion capture data to improve accuracy and compensate for a known misalignment between the human and exoskeleton joint axes [52]. Positions of the reflective markers were extracted and labeled using the automatic identification models (AIM) feature provided by the QTM software. The methods used for inverse kinematics varied depending on whether the trials involved single or multiple joints. In single-joint trials, which involved strictly only elbow flexion/extension, the angle of the elbow was directly computed using the marker coordinates. Conversely, in multi-joint trials, although the main focus was on shoulder and elbow flexions and extensions, it was essential to consider the potential movements of the entire upper body to take into account its full dynamics. Consequently, a comprehensive inverse kinematic analysis was conducted using OpenSim. Finally, kinematic data were filtered (4^*th*^ order 3*Hz* low-pass Butterworth) to align the harmonic content of kinematic data to the EMG envelope and ease the extraction of time derivatives using numerical differentiation. Joint angles, velocities and accelerations are referred to as 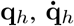 and 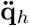.

#### 5) Human dynamics

Human dynamics were automatically computed from kinematics and anthropometric data previously introduced using Opensim. Classically, the motion of the human upper limb is described by the following dynamics, for *J* joints:

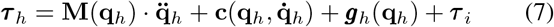

where, ***τ*** _*h*_ ∈ ℝ^*J*^ represents the torque exerted at the human’s joints, **M** ∈ ℝ^*J×J*^ denotes the mass matrix, **c** ∈ ℝ^*J*^ corresponds to the vector of Coriolis and centrifugal forces, ***g***_*h*_ ∈ ℝ^*J*^ signifies the vector of gravitational torques, and ***τ***_*i*_ represents the vector of human-exoskeleton interaction torques. By considering these factors, the equation provides a comprehensive representation of the complex dynamics involved in the motion of the human upper limb. Interaction torques are computed by first transforming the FT data from its original frame to the forearm’s frame and using the upper limb’s jacobian matrix **J**_*h*_(**q**_*h*_) so that

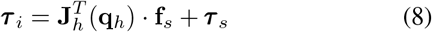

where **f**_*s*_ and ***τ*** _*s*_ are the forces and torques measured from the sensor underneath the orthosis.

#### 6) Resampling

All signals were resampled at a rate of 30*Hz* to reduce the number of samples and speed up the calibration of the models. All signals being 3*Hz* low-pass filtered, no information was lost.

### E. Analysis

As the present study aims to evaluate and compare EMG-to-torque models, it is necessary to properly define the criteria for selecting one model over another. In particular, for future implementations, the model accuracy and its computational cost (*i.e*., the calibration and evaluation durations) are essential performance criteria. The specific criteria to quantify these aspects are detailed below.

#### 1) Cross-validation

It is more conservative to provide different calibration and evaluation datasets to mitigate the risk of evaluating overfitted models. To that end, a K-fold cross-validation approach was employed. First, the dataset is divided into *k* folds. *k* iteration of calibration and evaluation are then performed. For each iteration, *k* − 1 folds are used for the calibration and the remaining fold is used for evaluation [53].

Due to the limited amount of data (10 trials per condition) and after testing various values of *k*, a two-fold division approach was used for this paper (see Fig. 3). For each participant and condition (single and multi-joint), the ten 30 s trials were randomly divided into two folds, each containing five trials. Therefore, for each model, condition, and participant combination, each of both resulting iterations yielded evaluation metrics. These two values were then averaged to obtain the scores of a model/condition/participant combination.

**Fig. 3:**
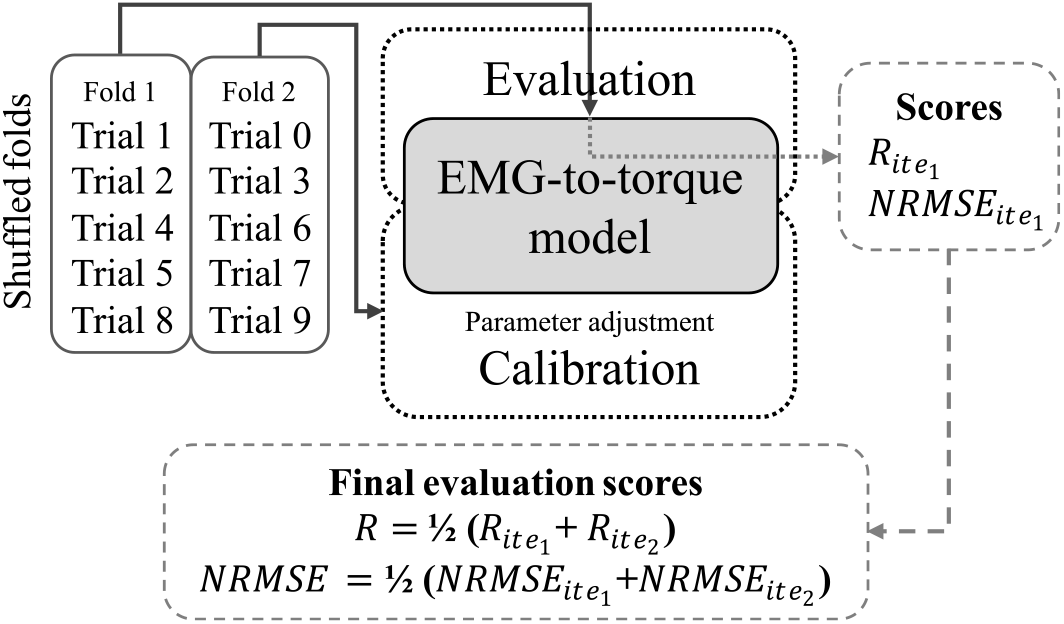
Cross-validation method. The data is split in two folds, one is used to perform the calibration and the other the evaluation. Roles are then reversed and the scores are averaged.

### 2) Performance metric

#### a) Joint torque estimation

The accuracy of the models was evaluated using two metrics: the normalized root mean square error (NRMSE) and Pearson’s coefficient of correlation (R). These metrics provide insights into the performance of calibrated models when estimating a joint torque in terms of data distance and error trends. The NRMSE is defined as follows

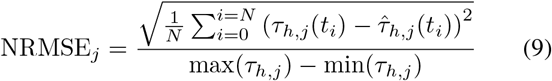

where *N* is the number of samples in the fold and *t*_*i*_ is the *i*^*th*^ time sample. The reference torque *τ*_*h*,*j*_ for the *j*^*th*^ joint is calculated from equation 7, while 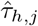 is an estimated torque obtained from the tested EMG-to-torque model. The NRMSE is normalized with respect to the amplitude of the reference torque, obtained using max(*τ*_*h*,*j*_) and min(*τ*_*h*,*j*_). Similarly, Pearson’s coefficient of correlation (*R*_*j*_) is computed as

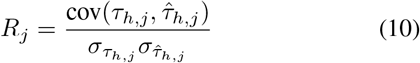

where cov(·, ·) denotes the covariance, and 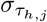 represents the standard deviation. The NRMSE and R values were compared during the cross-validation process to evaluate and compare the performance of the models.

### b) Computation efficiency

While EMG-to-torque model accuracy is undoubtedly the most critical feature for exoskeleton control, it is also crucial to be as computationally efficient as possible. Indeed, for future applications, it is necessary to induce short calibration times, which will influence the usability of the control on a daily basis with different users, and quick evaluation times, which will impact the controller’s responsiveness. As such, during cross-validation, calibration and evaluation durations were measured. The calibration duration was measured for each k-fold iteration across conditions and participants. The estimation duration was measured similarly and then divided by the number of samples to evaluate a feasible control loop period in a real-time application.

### F. Statistical analysis

The 2-fold cross-validation process resulted in two values for each criterion, which were then averaged, leaving one value per subject and model as data. *R* scores were normalized using the Fisher transform. The data was initially evaluated for its sphericity using Mauchly’s test. Subsequently, a repeated measures analysis of variance (rm-ANOVA) was conducted with the models as the within-subjects factor, and a Greenhouse-Geisser correction was applied if the sphericity was violated. The null hypothesis was that models produced identical scores. Model-to-model post-hoc comparisons were performed with a paired sample t-test and a Holm-Bonferroni correction. In the results, ± corresponds to the standard deviation (SD). Datapoints with a Z-score over 3 were considered to be outliers and not included in the statistical analysis.

## III. EMG-to-torque Models

The following section describes models using *K* muscles and *J* joints. Data matrices contain *N* samples.

### A. Linear mapping model

The multivariate linear regression (MVLR) model is a straightforward formulation used to compute joint torques by linearly combining muscle excitations [31]. This is achieved with a mapping matrix, denoted as **H**_𝓁_ ∈ ℝ^*J×K*^, which establishes the link between the estimated torques 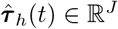 and the muscle excitations 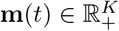. This relationship can be expressed as

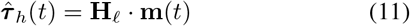

The model is calibrated by first defining the data matrices **T** ∈ ℝ^*J×N*^ and 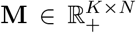, which contain torque and muscle excitation samples from the training dataset, respectively. The calibration process involves finding the mapping matrix **H**_𝓁_ that satisfies the equation:

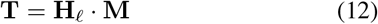

To achieve this, the *mvregress* function from Matlab (MathWorks, Natick, MA, US) is employed on **T** and **M** to determine the optimal values for **H**_𝓁_.

### B. Neuromusculoskeletal model

Neuromusculoskeletal (NMS) models are a combination of a neuromuscular Hill-type model [54] and a musculoskeletal representation of the body. A previously developed [50] and benchmarked [55] NMS model implemented on the OpenSim simulation software [51] was used in this study. Muscle activations are first derived from excitations via a nonlinear activation function [56] as follows

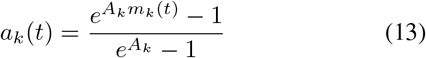

where *A*_*k*_, *a*_*k*_(*t*), and *m*_*k*_(*t*) are, respectively, the shape factor, activation and excitation of the *k*^*th*^ muscle. For a given muscle, the force can then be expressed as follows (see also [57]),

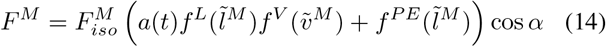

where 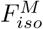 denotes the maximal isometric force, while 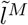 and 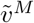 represent the normalized fiber length and velocity, respectively. The function 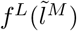 characterizes the forcelength relationship, describing how a muscle fiber’s maximum active force varies with its length. The normalized fiber length is given by 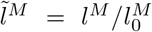, where 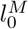 corresponds to the optimal fiber length at which 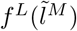 reaches its maximum value (*i.e*., 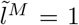). Similarly, 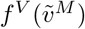 describes the force-velocity relationship, reflecting the changes in active force generation of the muscle with fiber velocity (*i.e*., *f* ^*V*^ (0) = 1 for isometric contractions), and 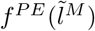 accounts for the passive elastic behavior of the fiber. Lastly, *α* denotes the pennation angle, which quantifies the angle between the axis of muscle force and the orientation of its fibers. This model incorporates a serial elastic element representing the tendon to ensure equilibrium between the muscle fiber and tendon forces and compute their lengths. This is captured by ensuring the following constraint during forward dynamics:

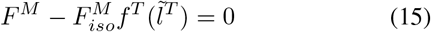

where 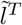 represents the normalized tendon length, and 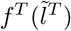 describes the force-length relationship of the tendon’s passive elastic behavior. The normalized tendon length is given by 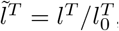, where 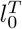 corresponds to the tendon slack length, such that 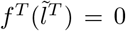 when 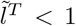. Equation 15 is solved at each time step during forward estimation of muscle force. Once the muscle force is obtained, the resulting estimated joint torques can then be derived as

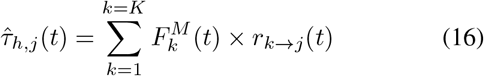

where *r*_*k*→*j*_(*t*) is the moment arm of *k*^*th*^ muscle on the *j*^*th*^ joint. During estimation and calibration, both the muscle force and the moment arm [58] are computed via OpenSim. When a muscle is not acting on a joint (*i.e*., brachioradialis on shoulder flexion), *r*_*k*→*j*_ is null.

The model is calibrated by first scaling the musculoskeletal component using anthropometric data from motion capture, adjusting the length and width of each limb. Then, the muscle parameters are optimized using a single objective genetic algorithm [29], [59] with a stalling criterion of 20 generations, 16 chromosomes per generation, and a maximum of 1000 generations. Crossover fraction and mutation rate were determined with a parametric study showing that a value of 0.7 for each provided the best calibration. The cost function is defined as the quadratic mean of the estimation mean square error of each joint:

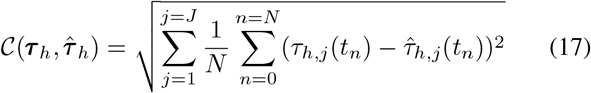

Five parameters are optimized for each muscle: maximal isometric force 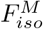, optimal fiber length 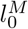, tendon slack length 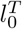, pennation angle at optimal fiber length 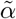 and shape factor *A*.

### C. Synergy-based model

Synergies are described as a *coherent activation, in space or time, of a group of muscles* [60]. A variety of synergy decompositions, including spatial, temporal, spatiotemporal, or space-by-time, can be extracted from a set of muscle excitations depending on the specific analysis or application [61]. However, since our dataset consists of randomly generated trajectories without specific gait or periodic movements, we do not anticipate the emergence of temporal synergies. Therefore, we have decided to focus on a spatial synergy model for our analysis. Considering the number of spatial synergies as *S*, the muscle excitation data matrix 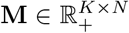 can be factorized into two nonnegative matrices as follows

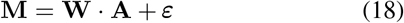

with 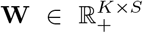 the synergy weight matrix, mapping a synergy activation to a group of muscles, and 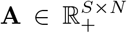 the synergies activation matrix, containing the corresponding synergy activation for each sample of the data. ***ε*** ∈ ℝ^*K×N*^ is the muscle excitation residual matrix that contains the residual error resulting from the factorization. After factorization, the synergy-based EMG-to-torque models are formulated similarly to the MVLR,

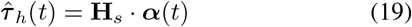

where **H**_*s*_ ∈ ℝ^*J×S*^ is the synergy mapping matrix, which maps a synergy to a joint, and 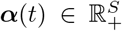 is the synergy activation vector (*i.e*., one of the columns of **A**). In this framework, ***α***(*t*) is not measured directly and must be calculated from the previously factorized synergy weight matrix:

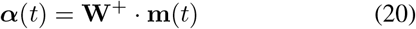

with •^+^ the pseudo-inverse operator. Therefore, the final formulation of the synergy-based model is

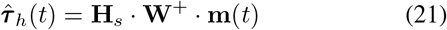

The model is calibrated in several steps. First, the muscle excitation matrix **M** from the training data (with a corresponding torque matrix **T**) is factorized via Matlab’s NNMF function to obtain the synergy weights and activation matrices **W** and **A**. Then, the calibration problem corresponds to finding **H**_*s*_ such that

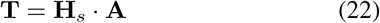

Similarly to the MVLR model, this is done by using Matlab’s *mvregress* function on **T** and **A**. In this paper, synergy-based models are abbreviated SYN; if the number of synergies is relevant in the context, the model is abbreviated SYNX, with X replaced by the number of synergies. To ensure at least one synergy per degree of freedom and direction, a minimum of 2 and 4 synergies were used for single and multi-joint conditions, respectively.

### D. Nonlinear mapping model

While the MVLR model is easy to formulate and calibrate, its main limitation stems from its inability to consider nonlinear muscle activation effects. To address this, we incorporate the shape factor, as described for the neuromusculoskeletal model, to obtain a nonlinear mapping (NLMap). The model formulation thus becomes:

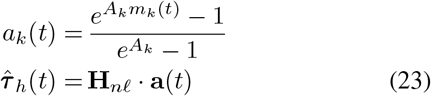

where *A*_*k*_, *m*_*k*_ and *a*_*k*_ are the corresponding shape factor, excitation, and nonlinear activation of the *k*^*th*^ muscle, and **H**_*n*𝓁_ ∈ ℝ^*J×K*^ the nonlinear mapping matrix. **a** is the activation vector.

The model is calibrated iteratively using a single-objective genetic algorithm with multiple steps in each iteration. Initially, the genetic algorithm generates a set of shape factors as a chromosome. These parameters are then used to compute the nonlinear muscle activations of the training dataset, which are compiled into an activation matrix **A**. Next, the mapping matrix is computed using the torque data matrix **T** through the *mvregress* function, solving the matrix form of equation 23. Since the mapping problem has a unique solution, a specific set of shape factors corresponds to a unique mapping, and hence, the matrix coefficients are not part of the chromosome. The final score of the chromosome is then computed with the cost function described in equation 17, and the best chromosome is determined through crossover and mutation operations in an iterative way as part of a typical genetic algorithm, with a 0.8 crossover rate and 200 chromosomes per generation.

## IV. Results

### A. Trajectory generation

The trajectory generation framework exhibited Gaussian-like distributions for elbow positions during single-joint and multi-joint conditions. The central tendency of elbow movements was observed around the 90^°^ position, with the majority falling within the 45^°^ to 135^°^ range (see Fig. 4). Shoulder positions were concentrated in the 0^°^ to 90^°^ range. The angular distributions observed during daily living tasks by Haverstock et al. [62] and Chapman et al. [63] are consistent with those produced by the framework. This shows that the framework can generate calibration trajectories that match expected use cases. During single-joint tasks, reference elbow torques were uniformly distributed within the −10 *N.m* to 20 *N.m* range. A Gaussian-like distribution emerged during multi-joint tasks, mainly featuring positive elbow torques ranging from −5 *N.m* to 20 *N.m* and shoulder torques spanning −10 *N.m* to 30 *N.m*.

**Fig. 4:**
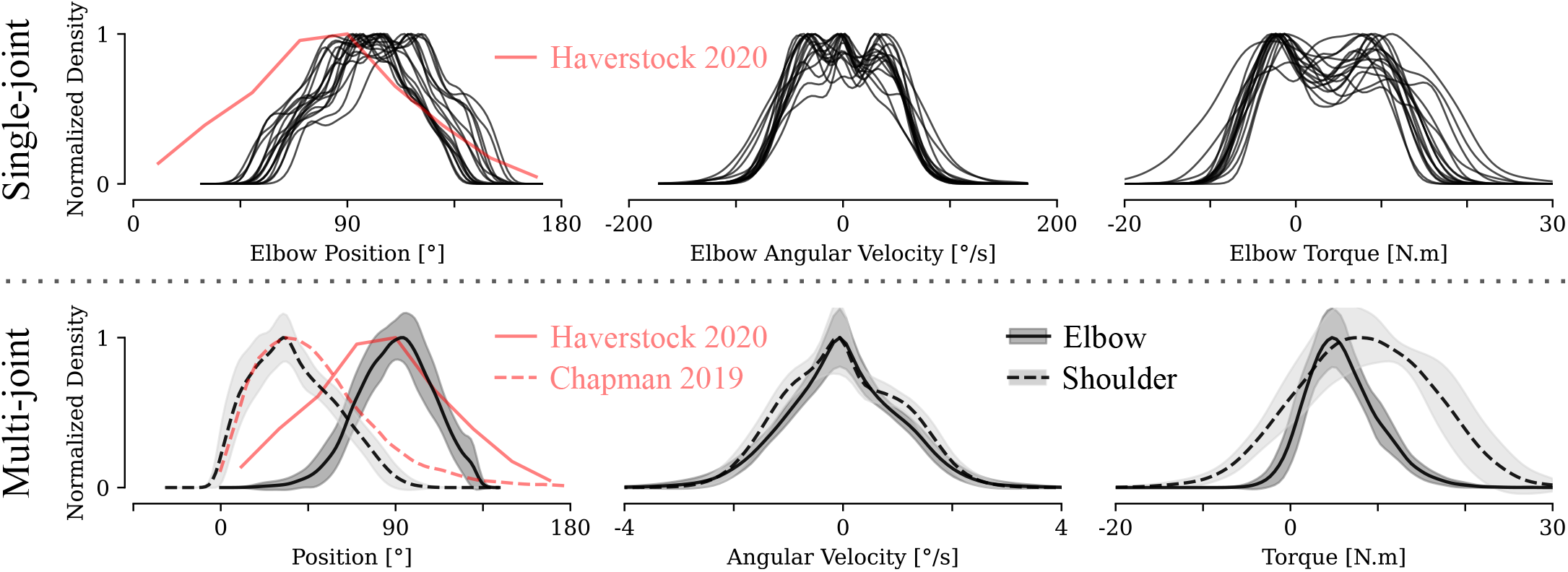
Generated trajectory kinematics and reference torque distributions. For the single joint condition, each line represents a participant. For the multi-joint condition, lines and fills represent the mean and standard deviation of the data distribution across all participants. Solid and dashed lines represent the elbow and shoulder joints, respectively. A 0^°^ angle corresponds to a fully extended elbow and no shoulder elevation. Areas around the lines correspond to the standard deviation between participants. Light red lines represent reference positions observed during daily living tasks for the elbow [62] and the shoulder [63].

### B. Torque estimation

#### 1) Single-joint

For the estimation of single-joint torque, the repeated measures ANOVA exhibited a notable influence of the chosen model on the NRMS error and the correlation coefficient (*p <* 0.001). Figure 5 presents a comprehensive summary of the performance outcomes. Specifically, the NLMap and NMS models attained NRMS errors of 4.66 ± 0.54% and 5.93 ± 0.96%, respectively. In contrast, the MVLR had a score of 7.11 ± 1.17%, and SYN models ranged from 7.31 ± 1.28% (SYN5) to 8.46 ± 2.37% (SYN2). Similarly, the correlation coefficient for the NLMap and NMS models were 0.971±0.011 and 0.958±0.013, respectively, while the MVLR was 0.942±0.019 and SYN models exhibited a range spanning from 0.943±0.016 (SYN6) to 0.918±0.0418 (SYN2). Overall, the NLMap model proved to be significantly better in terms of NRMSE and R (*p <* 0.001) than any other model with a strong effect (Cohen’s *d >* 1.65). The NMS model was also significantly better than MVLR and SYN models for both criteria (*p <* 0.05) with a strong effect (Cohen’s *d >* 1). Figure 6 visually compares correlations between the NLMap and MVLR models. The S-shaped correlation curve of the MVLR model indicates the presence of nonlinear residual errors. However, incorporating the shape factor term in the NLMap model contributes to their mitigation, resulting in a straighter curve that aligns more closely with the ideal 45^°^ line representing fully correlated torques.

**Fig. 5:**
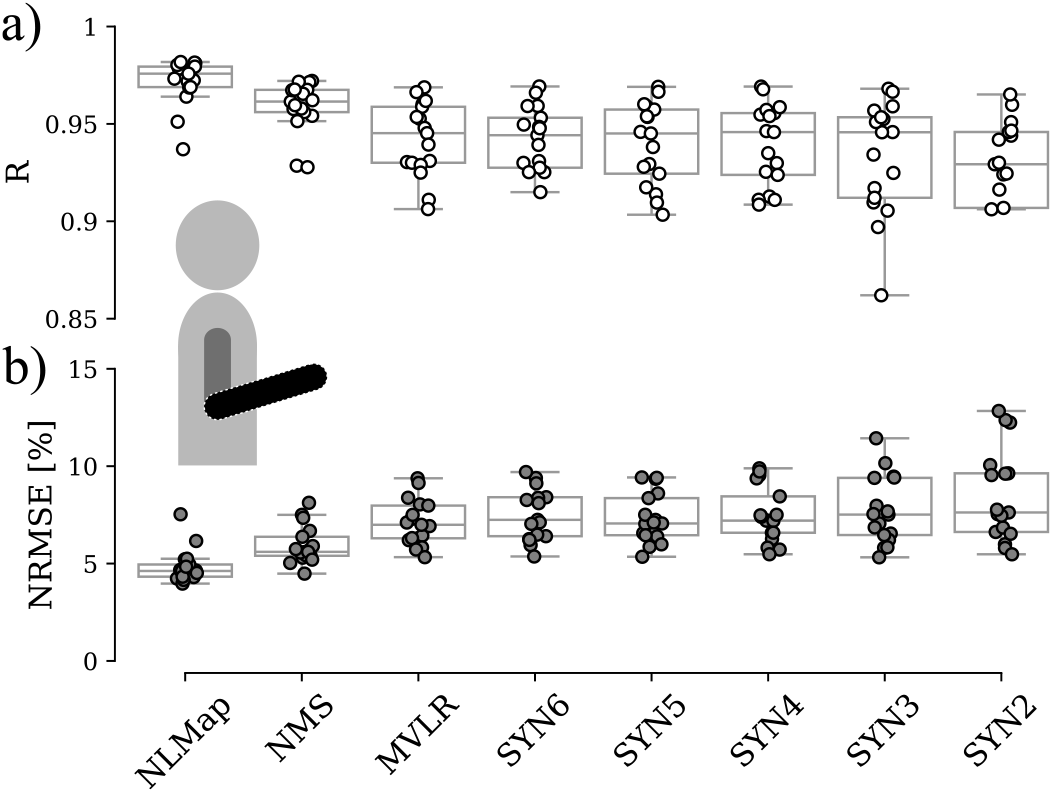
Single-joint model accuracy. a) Pearson’s correlation coefficient b) Normalized root mean square error

**Fig. 6:**
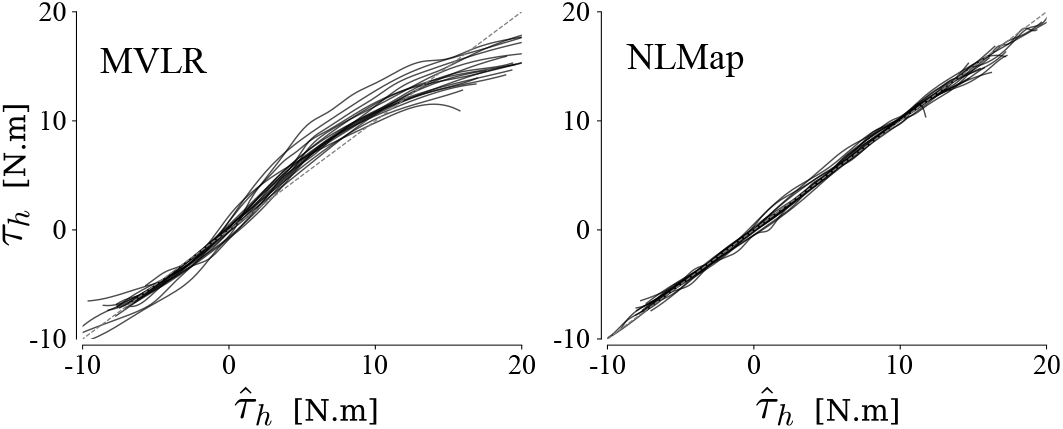
Effect of mapping nonlinearization on Elbow torques. Comparison between MVLR and NLMap of reference (*τ*_*h*_) to estimated 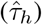 torques correlations. Each line corresponds to one participant and represents the smoothed data from the concatenated ten 30*s* trials.

#### 2) Multi-joints

The repeated measures ANOVA showed that model selection significantly affected the NRMS error and correlation coefficient for the shoulder and elbow joints (*p <* 0.01 and *p <* 0.001, respectively). The performance results are presented in Figure 8 for both joints. An illustration of the torque reconstruction capabilities of the NLMap mode is shown in Figure 7.

**Fig. 7:**
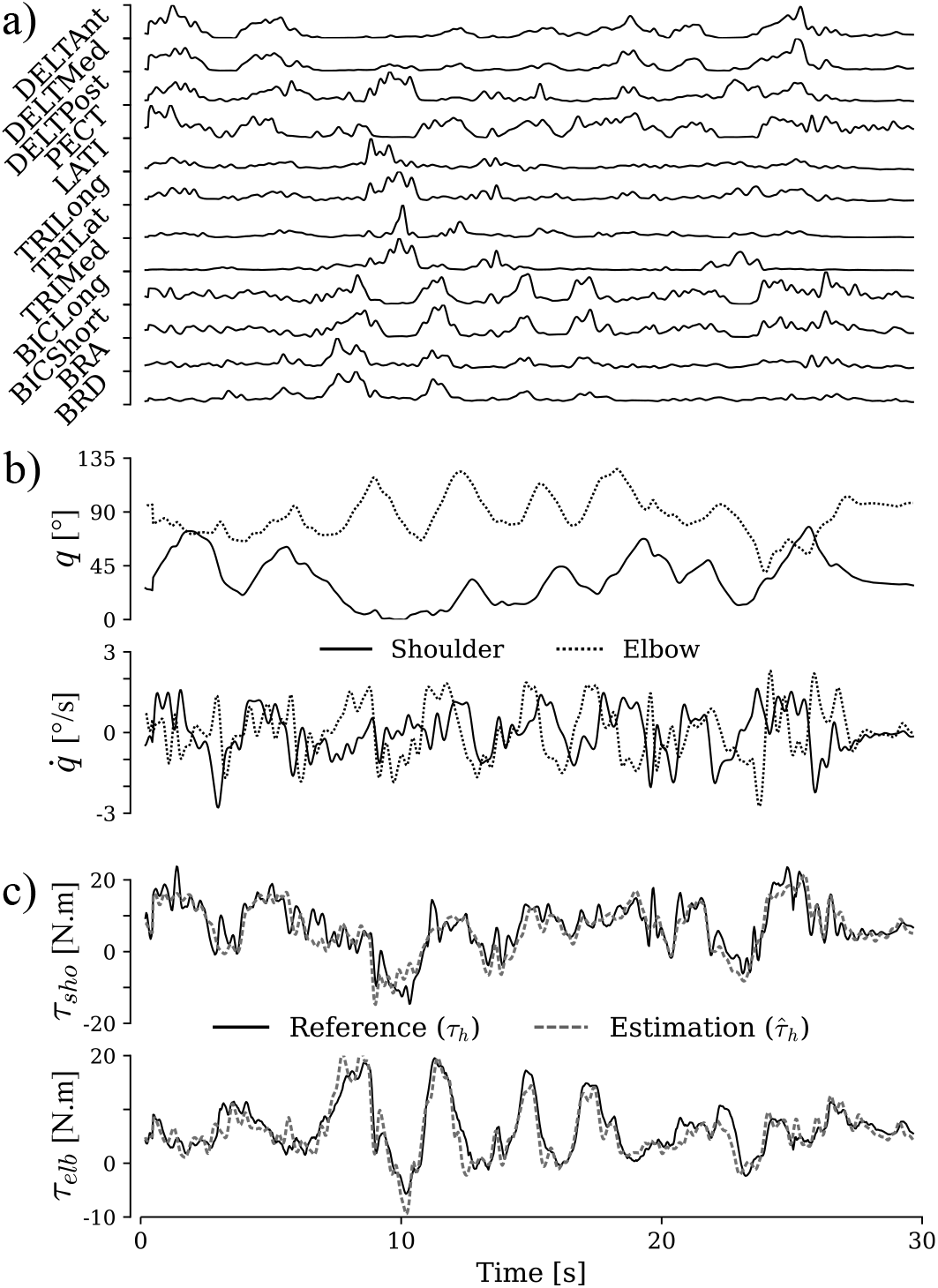
Multi-joint torque estimation with the NLMap model for a representative participant. a) Processed EMG signals. b) Angular position and velocity of the Shoulder (solid) and Elbow (dotted). c) Shoulder (*τ*_*sho*_) and Elbow (*τ*_*elb*_) reference and estimated torques. Reference and estimated torques are respectively represented using a solid and a dashed line.

**Fig. 8:**
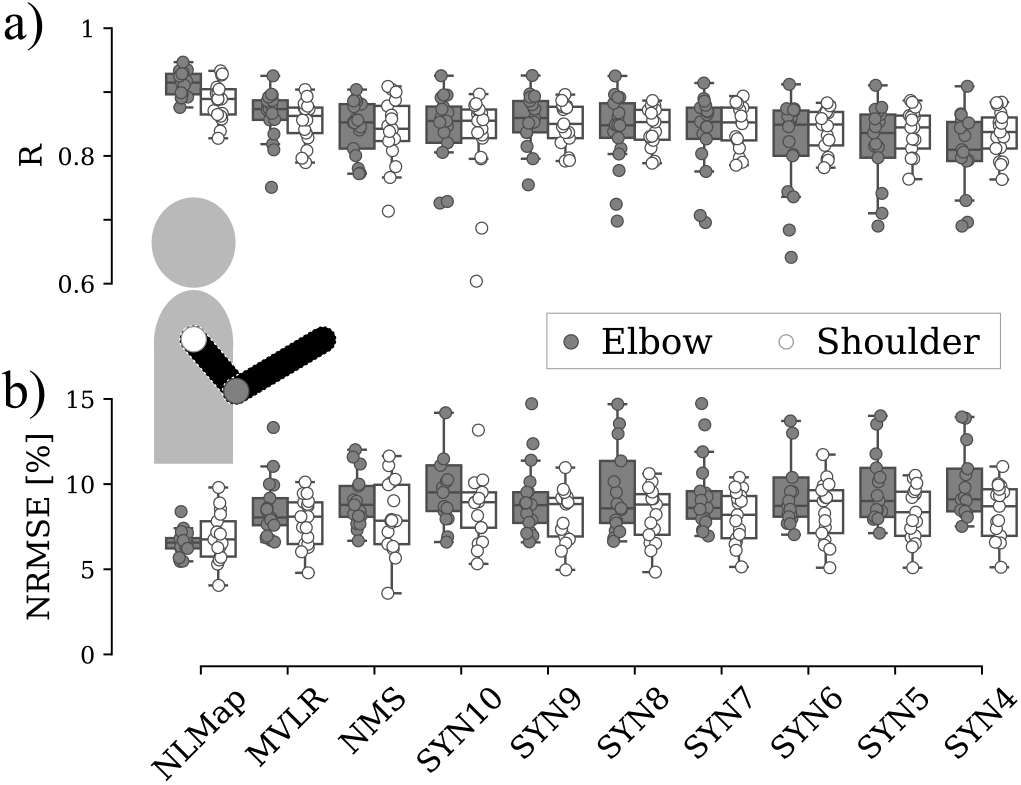
Multi-joint model accuracy. In both panels, data for the elbow joint is represented in gray, and data for the shoulder joint in white. a) Pearson’s correlation coefficient b) Normalized root mean square error

### a) Elbow

NLMap and NMS models obtained NRMS errors of 6.58 ± 0.73% and 8.97 ± 1.51%. The MVLR model had an error of 8.54 ± 1.75%, while SYN models ranged from 9.07 ± 2.12% (SYN9) to 10.24 ± 2.72% (SYN4). Likewise, NLMap and NMS models, respectively, obtained a correlation score of 0.913 ± 0.019 and 0.844 ± 0.043, while MVLR was 0.863 ± 0.041 and SYN ranged from 0.857 ± 0.042 (SYN9) to 0.798 ± 0.082 (SYN4). Overall, the NLMap model proved to be significantly (*p <* 0.01) better in terms of NRMSE and R than any other model with a strong effect (Cohen’s *d >* 1.45). On the other hand, the NMS model did not show significant differences from the MVLR model.

### b) Shoulder

The NRMS error for the NLMap and NMS models were 6.87 ± 1.4% and 8.0 ± 2.1%, respectively. The MVLR model had an error of 7.79±1.4%, while SYN models exhibited a range from 8.08 ± 1.5% (SYN7) to 8.47 ± 1.9% (SYN10). Similarly, the correlation scores for the NLMap and NMS models were 0.885 ± 0.030 and 0.840 ± 0.052, while MVLR was 0.855 ± 0.033 and SYN models ranged from 0.849 ± 0.034 (SYN9) to 0.828 ± 0.075 (SYN10). In terms of NRMSE, the NLMap model demonstrated statistically significant superiority (*p <* 0.05) with a moderate to strong effect (Cohen’s *d >* 0.62). Regarding the correlation coefficient, statistically significant differences (*p <* 0.01) were found between the NLMap and the other models. In particular, no significant differences were observed between the NMS and MVLR models.

### C. Computation efficiency

Duration results are shown in Table I. For linear models such as MVLR and SYN, the calibration time ranged from 1*s* to 2.5*s* in both conditions, while the NLMap model took an average of 22*s* to 45*s* to calibrate, depending on the condition. Finally, NMS models took the most time on average, ranging from 20 minutes in the single-joint condition to one hour in the multi-joint condition. Similarly, linear models were the fastest to estimate in a real-time setup, with an estimation time of less than a microsecond. The NLMap model took longer, with an average of a tenth of a millisecond. The NMS model was the longest, with an estimation time of 10.8*ms* for the single-joint condition and 15.6*ms* for the multi-joint condition.

**TABLE I:** Calibration and estimation duration. Calibration duration is the time necessary to perform the complete calibration of a model. Estimation duration is the time necessary to compute one estimation time step. The results with a value of *<* 10^*−*3^ indicate that the estimation duration was less than the resolution of the measurement.

| Single-joint | NLMap | NMS | MVLR | SYN6 | SYN2 |
| --- | --- | --- | --- | --- | --- |
| Calibration [s] | 22.1 | 1216 | 1.096 | 2.02 | 1.37 |
| SD [s] | 0.27 | 538 | 0.33 | 0.32 | 0.5 |
| Estimation [ms] | 0.08 | 10.8 | $< 10^{-3}$ | $< 10^{-3}$ | $< 10^{-3}$ |
| SD [ms] | 0.003 | 0.84 | $< 10^{-3}$ | $< 10^{-3}$ | $< 10^{-3}$ |
| Multi joints | NLMap | NMS | MVLR | SYN10 | SYN4 |
| Calibration [s] | 44.7 | 3662.14 | 1.49 | 2.45 | 1.89 |
| SD [s] | 2.6 | 1200 | 0.13 | 0.22 | 0.22 |
| Estimation [ms] | 0.1 | 15.6 | $< 10^{-3}$ | $< 10^{-3}$ | $< 10^{-3}$ |
| SD [ms] | 0.005 | 5.7 | $< 10^{-3}$ | $< 10^{-3}$ | $< 10^{-3}$ |

## V. Discussion

The present paper introduced a general framework for the *in situ* calibration and evaluation of EMG-to-torque models to enable predictive exoskeleton control to assist arbitrary human movements. We outlined a set of common models, their ontology, and the associated calibration technique. We also proposed a novel nonlinear mapping model designed to be simple in its formulation, with low calibration and estimation duration, yet able to consider nonlinear effects. We generated random, non-periodic trajectories, enabling us to test these models in various kinematic and dynamic situations. We investigated the difficulty of predicting torques with different degrees of freedom by testing single-joint and multi-joint strategies, which has yet to be addressed in the literature concerning arbitrary motions [9]. Finally, we evaluated the models using a cross-validation approach to prevent biases related to overfitting.

Creating a task to generate data with a wide range of kinematic and dynamic situations is essential to obtain a general calibration of EMG-to-torque models. While recent studies have used predefined or periodic movements [28], [64], the proposed framework was able to generate arbitrary random trajectories. Although the experimental setup prevented full coverage of the elbow positions, the distributions of angular positions were representative of daily tasks [62], [63]. This suggests that the presented framework is able to produce a model calibration that is especially valid in a workspace covering most daily activities and use cases.

Assessment of EMG-to-torque models necessitates the use of specific metrics for accuracy evaluation. This study utilizes Pearson’s correlation coefficient and the normalized root mean square error. While both metrics lead to similar conclusions about the models, their implications differ. Pearson’s correlation coefficient measures the linear relationship between the predicted and actual torques, inherently missing any nonlinear relationships. As depicted in figure 6, the MVLR model exhibited a nonlinear correlation with the reference torques, prompting the development of the NLMap model with a shape factor adjustment to enhance linear correlation. Conversely, the NRMSE measures the normalized difference between the estimated and actual torques, facilitating comparisons across different subjects and studies. However, it is highly sensitive to peaks in the actual torques, which can compress the normalized error and obscure discrepancies in error magnitude between low and high torque conditions. To address this issue, evaluations were conducted on trajectories that avoided high torque peaks, as confirmed in figure 6, showing no error variation between high and low torque for the NLMap model.

A key issue when calibrating and using EMG-to-torque models is their accuracy. We found that the NLMap model was the most accurate for both single-joint and multi-joint conditions. Given its simplicity, the MVLR model also demonstrated remarkable accuracy and should not be ignored for potential applications. Moreover, the NLMap and MVLR models do not require kinematic data as input features, making their implementation easier and necessitating fewer sensor types. The NMS model was surprisingly underperforming: while it had a satisfactory accuracy for single-joint estimation, it did not perform better than the MVLR in the multi-joint condition. This is due to the wide variety and the number of internal parameters that must be fitted, making it difficult to find a global minimum during calibration. Thus, while the NMS model should theoretically have the best accuracy due to its knowledge-based nature, this same nature mitigates the practicability of its calibration for in-field applications. The SYN models did not offer any notable advantages compared to their counterparts. Although the decrease in dimensionality may provide insight into how the central nervous system utilizes muscle redundancy during different movements, this does not appear to enhance the accuracy of movement prediction. It is difficult to compare the accuracy of other models in the literature because the evaluation method differs. Generally, models are evaluated on a different limb, with a different metric, feature, and task, and more often than not, without any cross-validation procedure. However, Table II provides an overview of some model scores reported in the literature using the NMRSE metric. The NRMSE scores for single-joint and multi-joint conditions indicated that the accuracy of the estimation for a single joint (*i.e*. the elbow) may be reduced when a more complex movement (*e.g*., shoulder rotations) is included. This effect could be caused by various factors, such as bi-articular muscles and movement artifacts due to the EMG sensor cables.

**TABLE II:** Overview of NMRSE model evaluations. Results are reported “as is”, taking the best score showcased in each paper. These results must be compared cautiously as they are reported using various evaluation methods, limbs, movements, and features. KF, LSTM, CNN, and NMF, respectively, stand for Kalman Filter, Long-Short Term Memory, Convolutional Neural Network, and Non-negative Matrix Factorization.

| Ref | Type | Feature | Joint | NRMSE [%] |
| --- | --- | --- | --- | --- |
| - | NLMap | Torque | Elbow | 6.50 |
| - | NLMap | Torque | Shoulder (Flex/Ext) | 6.87 |
| [32] | NMS | Torque | Wrist (Flex/Ext) | 2.02 |
| [65] | LSTM | Torque | Hip (Flex/Ext) | 3.6 |
| [38] | KF | Torque | Elbow | 5.81 |
| [66] | CNN-LSTM | Torque | Shoulder (Flex/Ext) | 12.44 |
| [66] | CNN-LSTM | Torque | Elbow | 12.76 |
| [30] | NMS | Angle | Wrist & Elbow | 13 |
| [33] | NMF-NMS | Angle | Wrist (Pro/Sup) | 17 |

While we tested the models on a powerful computer, exoskeleton controllers are typically embedded systems with limited computing power. Thus, it is important to assess the computational complexity of the models. However, to our knowledge, this kind of analysis has not been done in the literature. We found that simpler models, such as MVLR and SYN, were quicker to calibrate and provided faster real-time torque estimation. The calibration process took less than three seconds, and the duration of the estimation was so short that it could not be accurately measured (*<* 10^*−*3^ *ms*). The NLMap model calibration process took less than one minute, a suitable wait time for most applications. The estimation duration was less than one millisecond, allowing for a 1*kHz* exoskeleton control frequency. On the contrary, the NMS model did not demonstrate the same level of performance. Calibration took anywhere from 20 minutes to an hour, depending on the condition, and the estimation duration was more than ten milliseconds. This is due to the number of parameters that need to be adjusted during calibration (five parameters per muscle, sixty in total for multi-joint condition). Moreover, two steps are necessary for the computation of this model: muscle equilibrium and moment arm computation. The muscle equilibrium step is an iterative process that must be repeated at each time step, and for each muscle to balance the lengths of the fibers and tendons, it therefore influences computation duration both during calibration and estimation. Because muscle paths and insertions are not adjusted, the time taken to calculate the moment arm does not significantly impact the calibration duration since it is only necessary to do it once. However, during real-time estimation, each muscle’s moment arm must be computed for each actuated degree of freedom, impacting its duration. Consequently, numerical methods have been developed to accelerate this process [67].

This study offers new insights into the estimation of joint torques from EMG signals. However, certain limitations restrict the scope of these findings. To begin with, this research did not include all existing models as it would not have been feasible: we compared models commonly encountered in the literature and developed our model with practicality in mind. However, we proposed an evaluation protocol that enables anyone to test their model against others. To that end, we provided a standardized dataset that can be used to test new models and compare them with our results [68]. Currently, replicating the visual feedback of the trajectory and the motion-capture-based kinematic measurements for in-field use is challenging. Nevertheless, this visual feedback can be modified to display on wearable devices like smartphones or virtual reality headsets, and the human’s angular positions can be matched with those of the exoskeleton. Conversely, even though sagittal movements encompass many potential applications of exoskeleton assistance, these results should be extended to include all the degrees of freedom of the upper limbs. Furthermore, research has shown that the EMG signal can be degraded due to changes in the electrode/skin impedance over time, affecting the accuracy of the model estimation [32]. Consequently, sweat and skin degradation over time could impede exoskeleton assistance in a workplace environment, potentially making it unsafe. Although current EMG sensors allow the measurement of the electrode/skin impedance, no model has yet been developed to consider it in the torque estimation. Also, muscle fatigue may emerge as a factor impacting torque estimation accuracy over time. Fatigued muscles typically exhibit a lower mean EMG frequency [69] that current models fail to consider. Finally, other nonideal factors such as electrode shift and inter-day differences are known to induce accuracy variations [70], but could not be considered in this paper.

We anticipate conducting further research based on the results of this study. First, we will explore EMG feature extraction and reducing the number of measured muscles to enhance the robustness and widespread use of EMG-to-torque models. Comparing various feature extraction methods and muscle sets will be essential for improving the information embedded in the signal. Several studies have examined feature selection for EMG-based movement classification [22], [23], [71], but to our knowledge, no comprehensive comparison of EMG features has been conducted for continuous movement regression. Also, minimizing the number of measured muscles can simplify the placement of EMG sensors, which is a time-consuming and specialized task. Second, using our dataset and evaluation protocol, we plan to investigate artificial neural network-based (ANN) models, including Long Short-Term Memory (LSTM) networks. ANN models have the potential to offer more accurate estimations while maintaining a short calibration [21], [33]. They can be adapted to incorporate multiple EMG features and other relevant factors, such as kinematics. Finally, we aim to assess the impact of the estimation error on exoskeleton assistance and user feedback, considering potential safety concerns related to unexpected feedback loops and system instabilities resulting from variable human behavior and signal degradation over time. Despite these challenges, real-time estimation of human torques holds significant promises for the development of versatile assistive control strategies.

