## Supplementary material (S1 - S2) for "EMG-to-torque models for exoskeleton assistance: a framework for the evaluation of *in situ* calibration"

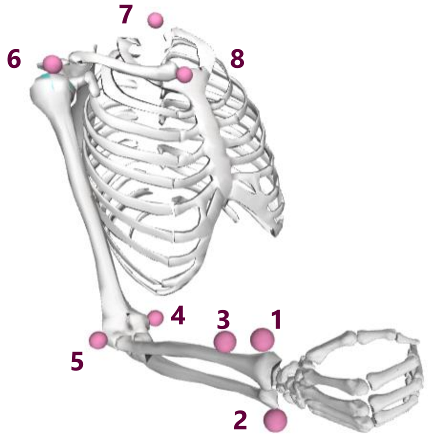


### S1. Motion capture measurements

#### a. Anthropometrics

Eight reflective markers were used during the anthropometric measurements. Seven of them were positioned on anatomical landmarks. An additional marker was employed to discriminate between pronation and supination.

| | |
|---|---|
| **Wrist**<br><br>1. Styloid process of radius<br>2. Styloid process of ulna<br>3. 4cm below styloid process of radius | 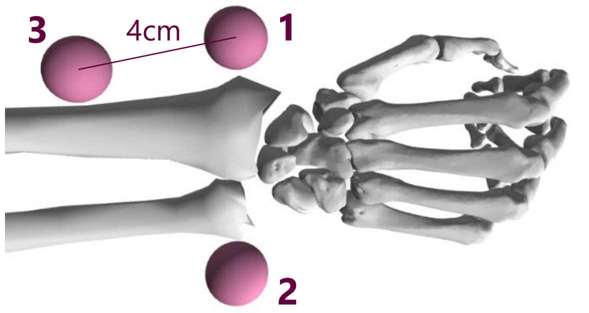 |
| **Elbow**<br><br>4. Lateral epicondyle<br>5. Medial epicondyle | 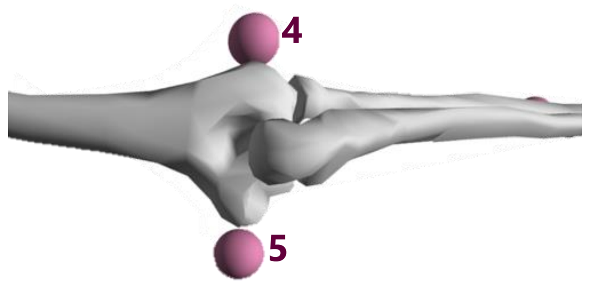 |
| **Upper torso**<br><br>6. Acromion<br>7. C7<br>8. Clavicle | 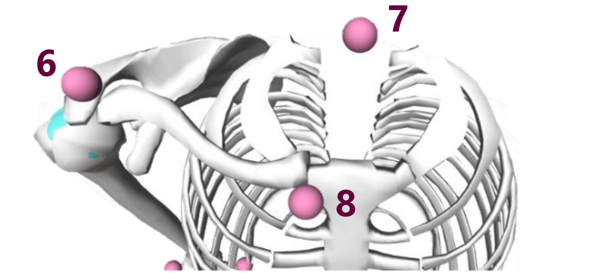 |

#### b. Experimental tasks

During the experimental tasks, additional markers were placed on the participant and the orthosis to enhance inverse kinematics and adapt to the presence of the exoskeleton.

The adjacent figure describes the reflective markers that were used after the anthropometric measurements were performed. The three orange markers (1, 2 and 8) were added on the front and back of the arm at the middle point, and on the distal part of the second metacarpal bone. The five blue markers (3-7) were directly positioned on the orthosis (see Figure 1); they allowed to track the position of the wrist. The blue markers were used to replace the styloid markers as they did not fit on the wrist while inside the orthosis. The styloid markers are denoted in green, signifying that they were removed for the task.

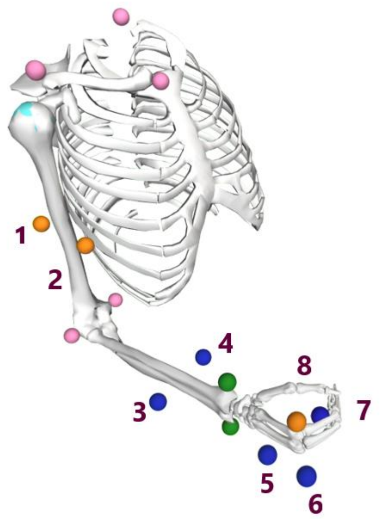

### S2. EMG Sensor placement

Twelve muscle heads were measured during this experiment:

- Anterior Deltoid
- Median Deltoid
- Posterior Deltoid
- Pectoralis Major
- Latissimus Dorsi
- Long Biceps
- Short Biceps
- Brachialis
- Brachioradialis
- Long Triceps
- Lateral Triceps
- Median Triceps

Sensors were placed according to SENIAM recommendations, and following illustration from the book by Perotto [1]:

| # | NAME | COMMENT | ILLUSTRATION |
|---|---|---|---|
| **1** | Anterior Deltoid | Arm along the body, supinated. | 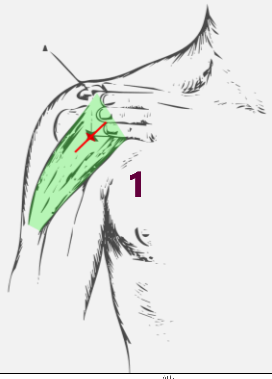 |
| **2** | Median Deltoid | Arm along the body, supinated. | 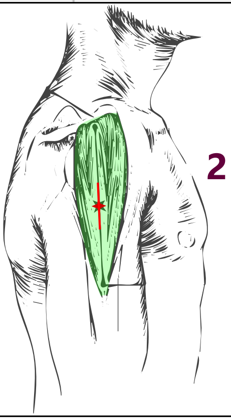 |
| **3** | Posterior Deltoid | Arm abducted. | 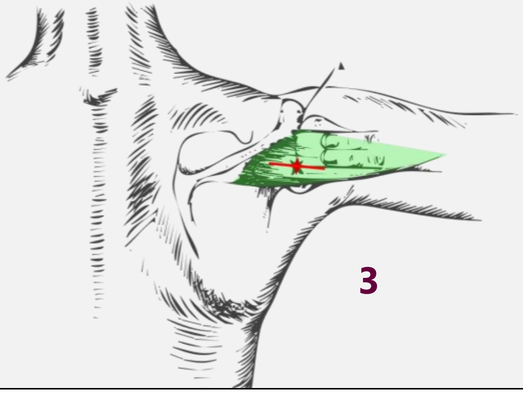 |

| | | | |
|---|---|---|---|
| **4** | Pectoralis Major | Clavicular head. | 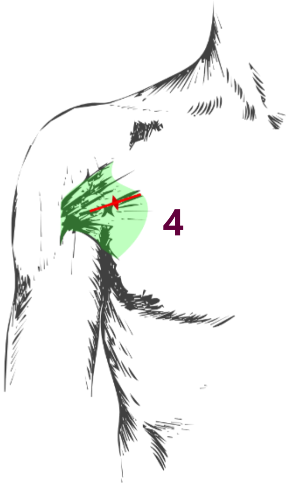 |
| **5** | Latissimus Dorsi | Arm along the body, supinated. | 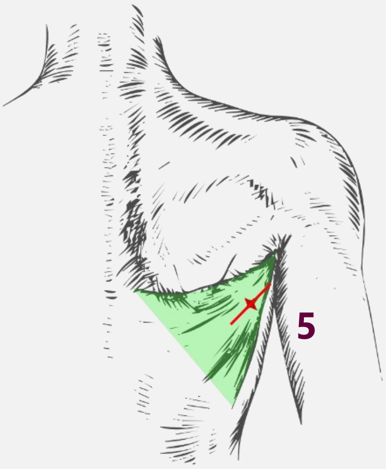 |
| **6** | Long Triceps | Arm abducted. | 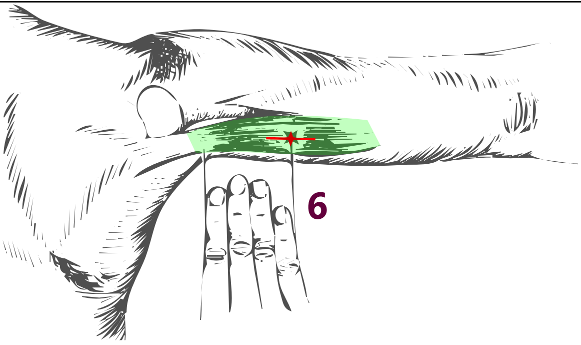 |

| | | | |
|---|---|---|---|
| **7** | Lateral Triceps | Arm along the bodu, pronated. | 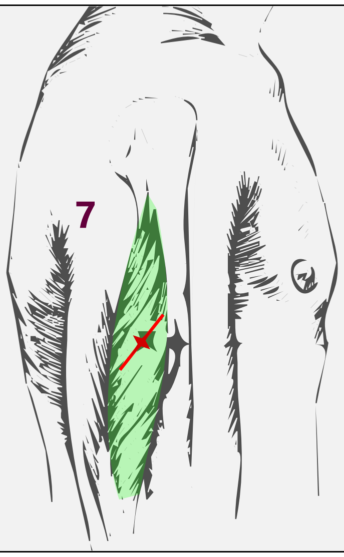 |
| **8** | Median Triceps | Arm abducted. | 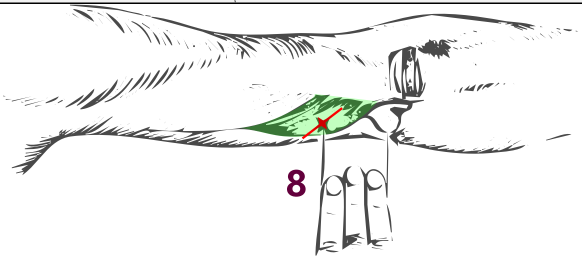 |
| **9** | Long Biceps | Exterior belly of the biceps. | 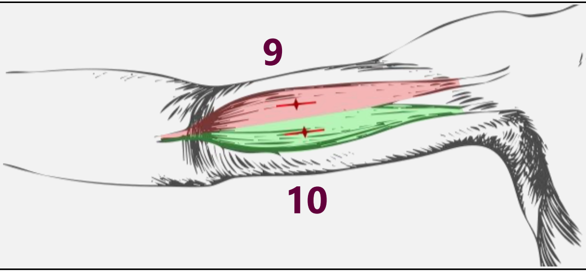 |
| **10** | Short Biceps | Interior belly of the biceps. | |
| **11** | Brachialis | Findable during elbow flexion. | 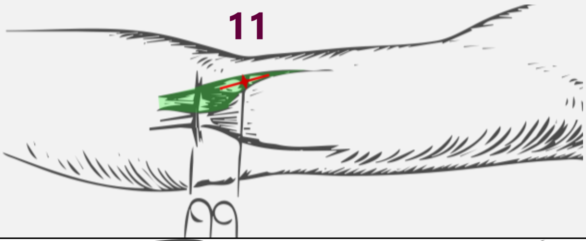 |
| **12** | Brachioradialis | Findable during elbow flexion. | 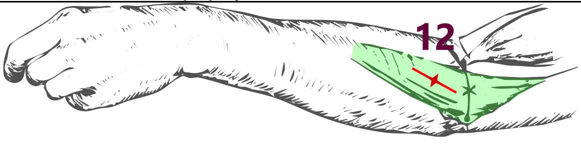 |

[1] Illustrations are adapted from "Anatomical guide for the electromyographer : the limbs and trunk." Aldo Perotto, 2011.

### S3. Single-joint task video

See the video attached to the paper. It shows 30s trial during the single-joint condition.

### S4. Multi-joint task video

See the video attached to the paper. It shows 30s trial during the multi-joint condition.
